# Human-derived NLS enhance the gene transfer efficiency of chitosan

**DOI:** 10.1101/2020.03.13.990390

**Authors:** Diogo B. Bitoque, Joana Morais, Ana V. Oliveira, Raquel L. Sequeira, Sofia M. Calado, Tiago M. Fortunato, Sónia Simão, Ana M. Rosa da Costa, Gabriela A. Silva

**Affiliations:** CEDOC, NOVA Medical School, Universidade Nova de Lisboa, Campo dos Mártires da Pátria, 130, 1169-056 Lisboa, Portugal; Algarve Chemistry Research Centre (CIQA), University of Algarve, Faro, Portugal; Centre for Biomedical Research (CBMR), University of Algarve, Campus Gambelas, 8005 Faro, Portugal

**Keywords:** gene therapy, chitosan, nuclear localization signals, IGFBP, HEK293T cells

## Abstract

Nuclear import is considered one of the major limitations for non-viral gene delivery systems and the incorporation of nuclear localization signals (NLS) that mediate nuclear intake can be used as a strategy to enhance internalization of exogenous DNA. In this work, human-derived endogenous NLS peptides based on Insulin Growth Factor Binding Proteins (IGFBP), namely IGFBP-3 and IGFBP-5, were tested for their ability to improve nuclear translocation of genetic material by non-viral vectors. Several strategies were tested to determine their effect on chitosan mediated transfection efficiency: co-administration with polyplexes, co-complexation at the time of polyplex formation, and covalent ligation to chitosan. Our results show that co-complexation and covalent ligation of human-derived NLS peptides to chitosan polyplexes yields a 2-fold increase in transfection efficiency.

These results indicate that the integration of IGFBP-NLS peptides into polyplexes has potential as a strategy to enhance the efficiency of non-viral vectors.

## Introduction

Gene therapy entails the transfer of therapeutic genetic material into cells where the production of the encoded protein will occur to treat or prevent a disease, by replacement of a missing or defective gene [1-4]. Gene therapy is a promising strategy for the treatment of many genetic and acquired diseases [2, 5]. Therefore, its success requires a gene delivery system with minimal toxicity, capable of protecting its load from degradation until it reaches its target, and allowing prolonged and stable gene expression [3, 6].

Non-viral gene delivery systems based on polyplexes have been considered a safe alternative to viral vectors due to their low toxicity, lack of significant immune response, ability to be administered repeatedly, low cost, and capacity to package large plasmids [7]. Nevertheless, the obstacles at the intracellular level [8] hinder transfection efficiency and are the major challenge for polymer-based gene therapy. For a successful gene delivery, polyplexes must efficiently enter the cell and traffic through the intracellular millieu towards the nucleus, overcoming biological barriers such as plasma, endosomal, and nuclear membranes [9]. We have previously explored the gene delivery properties of chitosan and hyaluronic acid [10-14] and confirmed the cause of the low transfection efficiency to be the inability of the polyplex/DNA load to enter the nucleus. To overcome these obstacles, we have explored chemical modification of the vectors [10, 12] as a strategy to increase the gene transfer efficiency. In this work, we evaluate the efficiency of transfection of our chitosan-based vectors after the incorporation of nuclear localization signals (NLS).

NLS are cationic peptide sequences that consist of either one or two stretches of basic amino acids of arginine/lysine and recognized by importins that direct their transport into the nucleus [4, 15]. NLS bind either directly to importin-β or to the adapter protein importin-α which, in turn, binds to importin-β and forms a complex. The resulting complex binds to the nuclear pore complex (NPC), by association with its cytoplasmic filaments, and is translocated through the pore to the nucleus. Finally, the complex dissociates and importins are recycled back to the cytoplasm, becoming available for a next import cycle [8, 16-18].

The use of NLS for non-viral gene therapy has been investigated due to the inefficient transfer of DNA to the nucleus. This is particularly relevant for post-mitotic and quiescent cells, where in the absence of mitosis there is no disintegration of nuclear membrane and nuclear entrance can only occur through the nuclear pore [19-21]. Regarding NLS efficiency in the nuclear transport of DNA, it has been found to be influenced by both the size and type of DNA (linear or plasmid), the method used for the incorporation of NLS (covalent or non-covalent attachment of NLS to DNA or polymer), and type of polymer used [15, 22].

Insulin-like growth factor binding proteins (IGFBPs) are a family of six mammalian multifunctional proteins [23, 24], which are involved in the regulation and transport of insulin-like growth factors IGF-I and IGF-II [23, 25]. Site-specific mutagenesis has revealed that the C-terminal region of both IGFBP-3 and IGFBP-5 (Table I), contains a domain with strong homology with known NLS sequences [23, 24, 26]. Mutations in this sequence lead to a reduction of nuclear accumulation [26, 27], suggesting that this 18 amino acid-long region of IGFBP-3 and IGFBP-5 is essential and sufficient for nuclear uptake and accumulation of IGFBP-3 and IGFBP-5 in several cell lines [23, 24, 26].

**Table I.**
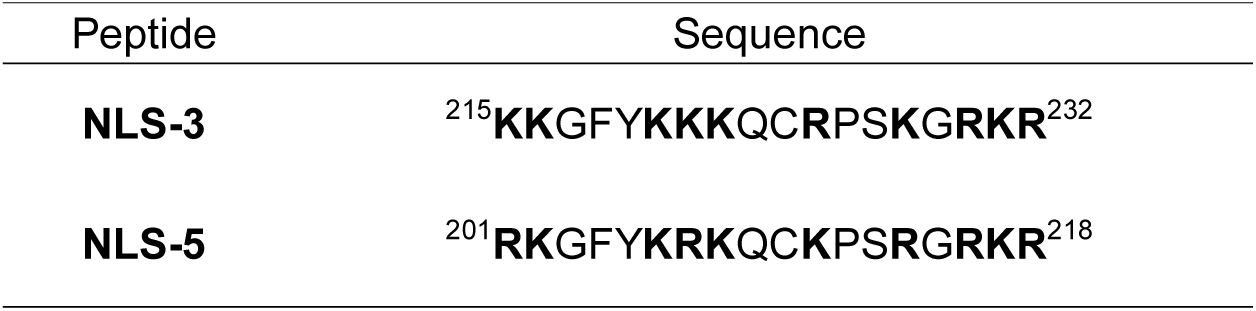
NLS sequences derived from the C-terminal region of IGFBP-3 (NLS-3) and IGFBP-5 (NLS-5); residues positively charged at physiological pH marked in bold.

These IGFBP peptides were already used to successfully deliver heterologous proteins, such as Glutathione *S*-transferase (GST) [28]; however, to the best of our knowledge, they have never been tested for gene delivery.

In this context, our aim was to incorporate NLS derived from IGFBPs into chitosan polyplexes to overcome the nuclear barrier and enhance nuclear internalization of DNA, thus increasing gene expression. We have tested three strategies to incorporate NLS into the formulations: co-administration of NLS and polyplexes, covalent ligation of NLS to chitosan, and co-complexation at time of polyplex formation. The polyplexes were extensively characterized regarding their size, polydispersity, charge, efficiency of DNA complexation, and cytotoxicity. We have found that the transfection efficiency of NLS-conjugated polyplexes, evaluated *in vitro* using HEK293T cells, was improved by two-fold.

## Materials and Methods

### 1. Materials

#### 1.1. Plasmids

A plasmid encoding the enhanced green fluorescent protein (eGFP) driven by the cytomegalovirus (CMV) promoter (pCMVeGFP) with the ampicillin resistance gene, was used for polyplex production. pCMVIGFBP-3 and pCMVIGFBP-5, driven by the CMV promoter were used to produce the NLS peptides derived from IGFBP-3 and IGFBP-5, respectively (Table I) containing a histidine tag (six residues) at the N-terminal.

The plasmids were amplified in Top10 *E. coli* bacteria (as described in section 2.1) and extracted using a Plasmid Maxi kit (Qiagen, Germany), according to the manufacturer’s instructions. Plasmids were later dissolved in TE buffer and their concentration was determined at 260 nm using a NanoDrop 2000c spectrophotometer (Thermo Scientific, USA).

#### 1.2. Polymers

Ultrapure chitosan (CS), with MW of 80 kDa and degree of deacetylation of 83%, was purchased from Novamatrix (CL 113, FMC BioPolymer AS, Norway). Polymer solutions of 1 mg/ml were prepared by dissolving the polymer in ddH_2_O, and the pH of the solutions adjusted to 5.5 with sodium hydroxide. All solutions were sterile filtered through a 0.2 μm filter.

All other reagents were acquired from Sigma-Aldrich (St. Louis, MO/USA).

#### 1.3. Cell culture

The HEK293T cell line was used for the transfection assays. Cells were cultured in Dulbecco’s modified Eagle’s medium (DMEM), containing 10% fetal bovine serum (FBS), 1% glutamine and 1% penicillin/streptomycin solution, and maintained at 37°C under a 5% CO_2_ atmosphere. Cell culture reagents were acquired from Sigma-Aldrich (St. Louis, MO/USA).

### 2. Methods

#### 2.1. Bacterial transformation

For bacterial transformation, *E. coli* TOP 10 bacteria were thawed and kept on ice and transformed by heat shock at 42°C for 90 seconds using 30 ng of plasmid (pCMVIGFBP-3 or pCMVIGFBP-5) Afterwards, 300 µl of SOC medium (98% tryptone, yeast extract and NaCl, 1% of Mg^2+^ and 1% of glucose) was added and the bacterial suspension was incubated at 37°C for 30 minutes under constant stirring (180 rpm). The transformed bacteria (100 µl) were spread in pre-warmed LB agar plates containing kanamycin (30 µg/ml) and incubated overnight at 37°C.

#### 2.2. NLS peptide production and purification

The IGFBP-derived NLS peptides (NLS-3 and NLS-5) were extracted from the bacteria using the B-Per 6xHis Fusion Protein Purification Kit (Thermo Scientific, USA), according to manufacturer’s instructions, and quantified by the Bradford method [29]. Thereafter, the NLS peptides were twice dialyzed (first against 10 mM HCl solution for 6 hours and then against ddH_2_O water for 12 hours) in dialysis tubing with a 2 kDa MW cut-off (Sigma-Aldrich, St. Louis, MO/USA). After dialysis, the NLS peptide solutions were frozen at −80°C and freeze-dried. Concentrations were further determined by spectrophotometry at 280 nm, using the molar extinction coefficient obtained from a calculation tool made available by Innovagen (http://pepcalc.com/).

#### 2.3. Polyplex preparation

To test non-viral formulations of chitosan/DNA polyplexes with NLS peptides, we have used different methodologies: co-administration of polyplexes and peptides at the time of transfection, covalent ligation of NLS peptides to chitosan, and co-complexation, by adding the NLS peptides at the time of chitosan-DNA polyplex formation. A summary of the produced formulations is shown in table II.

**Table II.** Polyplex formulations. CS – chitosan; N – nitrogen (representing amine groups); P – Phosphate (representing DNA). For formulation CS (chitosan), **Na**_**2**_**SO**_**4**_ was added to form the polyplexes.

| Formulation | NLS | Strategy |
| --- | --- | --- |
| CS0 | No | N : P = 15:1 |
| CS | No | N : P = 15:1<br>Co-administration of polyplexes and NLS peptides (NLS-3: 10, 25, 50, 100 and 150 $\mu\text{g}$; NLS-5: 10, 25 and 50 $\mu\text{g}$) |
| CS3(T) | NLS-3: 10, 50 and 100 $\mu\text{g}$ | N : P = 15:1<br>Co-complexation. NLS added to CS solution |
| CS5(T) | NLS-5: 10, 50 and 100 $\mu\text{g}$ | N : P = 15:1<br>Co-complexation. NLS added to CS solution |
| CS3(S) | NLS-3: 10, 50 and 100 $\mu\text{g}$ | N : P = 15:1<br>Co-complexation. NLS split between CS and DNA solutions |
| CS5(S) | NLS-5: 10, 50 and 100 $\mu\text{g}$ | N : P = 15:1<br>Co-complexation. NLS split between CS and DNA solutions |
| 15:1-CS3T | NLS-3: 100 $\mu\text{g}$ | N (CS+NLS) : P = 15:1<br>Co-complexation. NLS added to CS solution |
| 15:1-CS5T | NLS-5: 100 $\mu\text{g}$ | N (CS+NLS) : P = 15:1<br>Co-complexation. NLS added to CS solution |
| 15:1-CS3S | NLS-3: 100 $\mu\text{g}$ | N (CS+NLS) : P = 15:1<br>Co-complexation. NLS split between CS and DNA solutions |
| 15:1-CS5S | NLS-5: 100 $\mu\text{g}$ | N (CS+NLS) : P = 15:1<br>Co-complexation. NLS solution split between CS and DNA solutions |
| CSedac3 | NLS-3: 100 $\mu\text{g}$ | N (CS+NLS) : P = 15:1<br>NLS covalently linked to CS |
| CSedac5 | NLS-5: 100 $\mu\text{g}$ | N (CS+NLS) : P = 15:1<br>NLS covalently linked to CS |

To prepare chitosan-DNA polyplexes, DNA was diluted in 500 µl of a sodium sulfate solution (25 mM) and an equal volume of CS solution was preheated to 55°C for 5 min to prevent aggregation upon polyplex formation [30]. Both solutions were quickly mixed together, placed on ice for 30 min and stored at 4°C. Different amounts of NLS peptides (10, 25, 50, 100 and 150 µg for NLS-3 and 10, 25 and 50 µg for NLS-5) were co-administrated with this formulation (CS) at the time of the *in vitro* transfection. The N:P ratio (amines from CS and P from DNA) was 15:1.

To test if polyplexes were affected by the presence of salts, the DNA solution was not diluted into the sodium sulfate solution, and polyplexes were prepared by adding the DNA solution directly to the CS solution either in the presence (CS3 and CS5) or absence (CS0) of NLS peptides. The total amount of NLS peptides was either added to the CS solution [CS3 (T), CS5 (T)] or split into equal parts and added to each solution [CS3 (S), CS5 (S)] which were then quickly mixed as described above. Different amounts of NLS peptides were tested (10, 50 and 100 µg).

To test for differences in complexation efficiency, polyplexes were prepared considering the total amount of amine groups, adding those of NLS peptides and chitosan, at the same N:P ratio of 15:1. These formulations were prepared in the same way as described for the previous polyplexes (CS3 and CS5, S and T). For these formulations (15:1CS3 and 15:1CS5), a fixed peptide amount of 100 µg was chosen.

We have also covalently linked the NLS peptides to chitosan, by using a one and a half molar excess (relative to the carboxylic acid groups in the peptides) of *N*-(3-Dimethylaminopropyl)-*N*’-ethylcarbodiimide hydrochloride (EDAC) added to 100 µg of NLS peptides and stirred at 4°C for 24h. The mixture was then added to chitosan and polyplexes (formulations CSedac3 and CSedac5) prepared as described above and stored at 4°C.

#### 2.4. Polyplex characterization

Polyplex size measurements were performed with dilution in ddH_2_O water by dynamic light scattering (DLS), at 25°C with a detection angle of 173°, using a Zetasizer Nano ZS (Malvern Instruments, UK) and zeta potential (surface charge) measured with laser Doppler velocimetry.

#### 2.5. Polyplex complexation efficiency

DNA complexation efficiency by the polyplexes was assessed using a gel retardation assay. Agarose gels were prepared in TAE buffer (1% w/v) and visualized using GreenSafe® Premium (NZYtech, Portugal). Polyplex formulations were loaded in each well and electrophoresis was carried out for approximately 60 minutes at +90 mV, with samples visualized under an UV light.

#### 2.6. NLS influence on cell viability

To evaluate the effect of NLS-3 and NLS-5 peptides on HEK293T cell viability, an MTT [3-(4,5-dimethylthiazol-2-yl)-2,5-diphenyltetrazolium bromide] assay was performed. Cells were seeded in a 48-well plate at a density of 15 000 cells per well with 500 µl of complete DMEM (supplemented with FBS, pen-strep) and allowed to grow for 24 hours, at 37°C under a 5% CO_2_ atmosphere. Culture medium was then replaced by serum-free DMEM containing different amounts of NLS-3 (0, 10, 25, 50, 100 and 150 µg) and NLS-5 (0, 10, 25, and 50 µg) incubated for 5 hours, and then replaced by complete DMEM until analysis.

For the cytotoxicity evaluation, the cell culture medium was removed and replaced by 25 µl of MTT solution (5 mg/ml in PBS) and cells were further incubated for 4 hours. To solubilize the formed formazan crystals, 250 µl of a 0.04 N HCl in isopropanol solution was added to each well. After one hour, the absorbance was measured on a microplate reader (Tecan Infinite M200, USA) at 570 nm for formazan quantification, and at 630 nm for cellular debris. Non-treated cells (0 µg of NLS peptides) were used as positive control, and cells treated with latex extract were used as negative control for cell viability. Cell viability was calculated using the following equation:

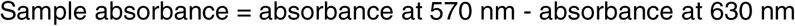

Cell viability (%) = (sample absorbance / positive control absorbance) × 100

All experiments were performed in triplicate.

#### 2.7. *In vitro* transfection assays

Cells were seeded at a density of 200 000 cells per well in 6-well culture plates with DMEM supplemented with FBS at 37°C under a 5% CO_2_ atmosphere. 24 hours after cell seeding, the formulations, prepared as described in section 2.3, were added to the cells, and followed the experimental setup described in the previous section. All formulations were added to the cells in a single step, except for the co-administration condition, where the polyplexes were added to the cells first, followed by different amounts of NLS-3 and NLS-5 peptides. Transfection efficiency was assessed qualitatively using a fluorescence microscope (Axiovert 40 CFL, Zeiss, Germany) for GFP expression at 48h and 72h post-transfection, and quantitatively by flow cytometry (FACSCalibur, BD Biosciences, USA). For the latter, cells were washed thrice and re-suspended in PBS for analysis of GFP expression, with a total of 50,000 events counted for each sample. All experiments were performed in triplicate.

#### 2.8. Statistical analysis

Statistical analysis was performed using Prism 6 (GraphPad Software), and statistical significance was analyzed by one-way ANOVA with multiple comparison tests or by an Unpaired t-test with Welch’s correction using a confidence interval of 95% and considering P<0.05 value as significant.

## Results & Discussion

### 3.1. Polyplex formulations display properties adequate for gene transfer

A positive surface charge is required for efficient cellular uptake since cell entry occurs by non-specific electrostatic interactions between the positively charged polyplexes and the negatively charged cell surface [31]. Parameters as N:P ratio, molecular weight, mixing technique, among others, have been widely investigated on CS-DNA complexes [30, 32-34], and described as factors that influence the binding affinity between chitosan and DNA, size and charge of polyplexes, cellular uptake and dissociation of DNA, and thus, transfection efficiency [35-37]. Based on our previous work [10-14], an amine to phosphate (N:P) ratio of 15:1 was chosen to promote the formation of positively charged polyplexes and avoid repulsion by the negative cell surface.

The preparation method of chitosan-DNA polyplexes was adapted from Mao *et al*, where polyplexes were prepared by solubilizing DNA in a Na_2_SO_4_ solution and mixing with a chitosan solution. Electrostatic interactions between positively charged amine groups (of chitosan) and negatively charged phosphate groups (of DNA) is the known driver for polyplex formation. This preparation method yielded homogeneous formulations (CS), with polydispersity below 0.3, mean size of 285.85 ± 56.50 nm and positively charged, with mean values of zeta potential of 15.45 ± 0.21 mV.

The effect of salt addition in the coacervation of polyelectrolytes (salting-out) is well known [38-40]. This effect has been correlated to the chaotropic/kosmotropic nature of the salt ions, contributing to phase separation by removal of the water layer around the dissolved macromolecules [30], to a greater or lesser extent, according to the Hofmeister series. However, it is not very likely that even strongly hydrated ions significantly influence water beyond their immediate solvation shells, and therefore the solute itself needs to be considered [41]. It is indeed more likely that this effect is a result of charge screening, where salt ions interact with the polyelectrolytes, providing local charge compensation for the charge imbalance and favoring more accessible polymer conformations, thus promoting chain entanglement [40]. Since Mao *et al*, observed that the presence of sodium sulfate within the range 2.5 - 25 mM did not significantly affect the mean size of CS/DNA polyplexes, in order to evaluate the effect of salt addition in the properties of the prepared polyplexes, this solution was completely removed from polyplex preparation [30].

Polyplexes prepared in the absence of sodium sulfate (formulation CS0) presented an increased mean size of 729.27 ± 116.77 nm and an increased positive charge of 27.17 ± 10.43 mV. This may be ascribed to stiffer chain conformations of both polyelectrolytes, due to the low ionic strength of the medium, creating obstacles to an effective chain entanglement and charge neutralization.

Using this preparation method, different amounts of NLS peptides (10, 50 and 100 μg) were co-complexed with polyplexes using two different approaches: the total amount of each NLS peptide added solely to the chitosan solution [formulations CS3 (T) and CS5 (T), with T for total] or divided into 2 equal parts and each part added to either the chitosan or DNA solutions [formulations CS3 (S) and CS5 (S), with S for split], as described in the methodology section. NLS-3 and −5 are 18 amino acid peptides bearing 10 positively charged residues (arginine and lysine) intercalated with neutral ones, a six histidine residues tag, which is partially protonated (∼78% at pH 5.5), the N-terminal ammonium group and the C-terminal carboxylate. With molecular weights of 3 kDa, their charge/mass ratio is approximately 5 × 10^−3^, practically the same as 83% deacetylated CS, with an average molar mass of 168 g/mol and a 0.83 positive charge per monomer. Therefore, the total N:P ratio in CS-3 and CS-5 formulations are approximately 15.5:1, 16.5:1 and 18:1 for the addition of 10, 50 and 100 μg, respectively.

These polyplexes were characterized regarding their size, polydispersity, and zeta potential, as shown in a representative figure (Figure 1) for S (Split for NLS split between chitosan and DNA solutions) and T (Total NLS added to chitosan) formulations. There seems to be a trend for size reduction with increasing NLS concentration, although not to a statistically significant extent, except for the two highest concentrations of NLS-5. A non-significant variation on zeta potential was also observed.

**Figure 1.**
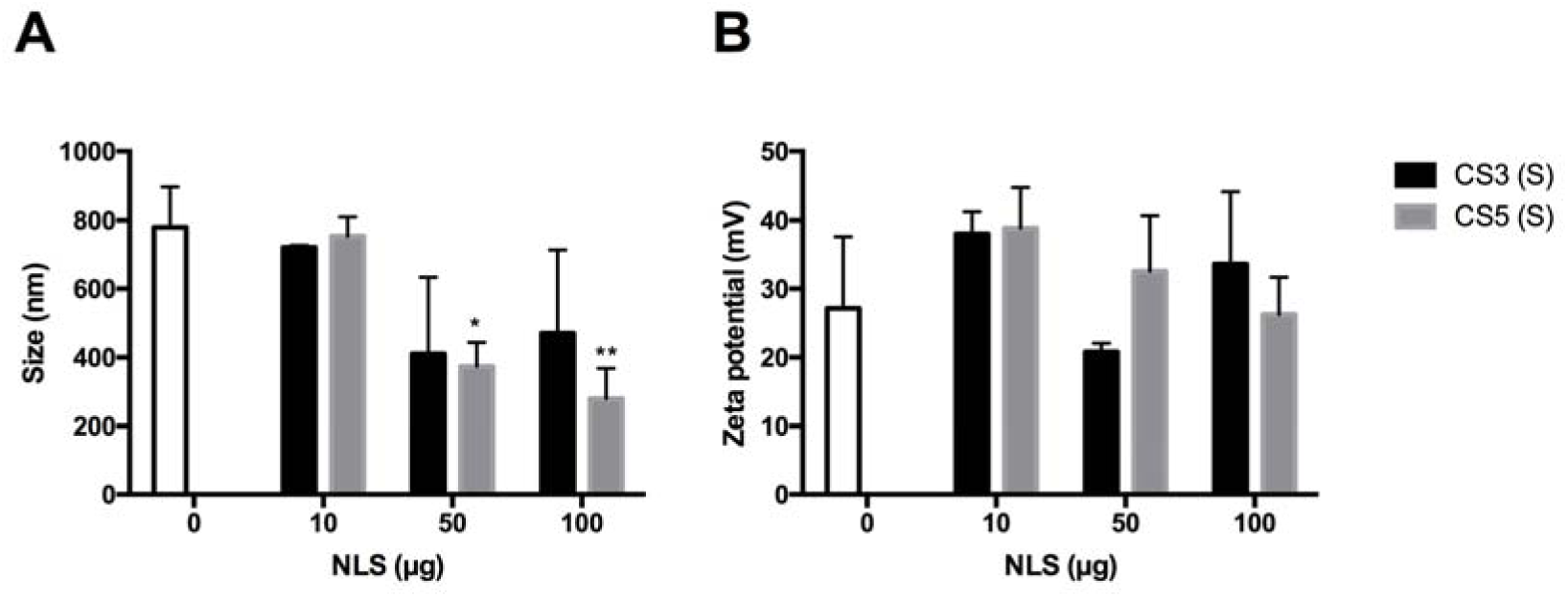
Physical characterization: Size (graph A) and Zeta Potential (graph B) of CS3 and CS5 polyplexes. The white bar represents the CS polyplexes without NLS. The graphs are representative for S and T formulations, as no differences were found between these formulations. Polydispersity was similar for all samples (below 0.4). Statistical differences, compared to polyplexes without NLS peptides (0) were calculated using Dunnett’s multiple comparisons test (N=3, error bars=SD; **p<0.01; *p<0.05).

Being relatively small molecules and containing positively charged groups dispersed throughout the backbone, NLS may have a bridging or reticulating effect during polyplex formation, leading to lower polyplex sizes. Another possibility is a competition between these molecules and CS for the negatively charged DNA chains, leading to the replacement of some CS chains in polyplexes and to more reduced sizes. The more significant effect of NLS-5 may be attributed to its higher arginine content: contrary to lysine, whose ammonium groups may be self-solvated by the carbonyl oxygen atoms of amides when sterically possible and therefore unavailable for salt bridging, the large guanidinium group of arginine, with an extensive delocalized charge, is less prone to self-solvation, hence more available for ion pairing [42]. Nevertheless, the addition of NLS seems to not affect zeta potential, which is consistent with both the above possibilities, as entrapment of NLS inside the polyplexes or replacement of CS chains by the peptide would not drastically change the surface charge.

Since the mixing method can influence the properties of polyplexes [35], we evaluated if adding the total amount of NLS peptides to the chitosan solution or adding it in equal amounts to either the chitosan or DNA solutions had an impact on polyplex formation. Because NLS, like CS, are positively charged, a competition between the two types of molecules for the DNA strands is expected. By partly adding the NLS to the DNA prior to polyplex formation, the inclusion of NLS in the formulations is guaranteed, either by formation of soluble complexes or even small coacervate particles [43], which will grow upon addition of CS and the remaining NLS. In this way, we expect to have some NLS chains at the surface of polyplexes, in case polyplexes approach the nucleus still intact, and also near the plasmid, in case polyplex degradation starts during lysosomal escape and only DNA reaches the nucleus. We have tested all conditions and selected a representative condition, corresponding to an NLS amount of 100 µg, as shown in Figure 2.

**Figure 2.**
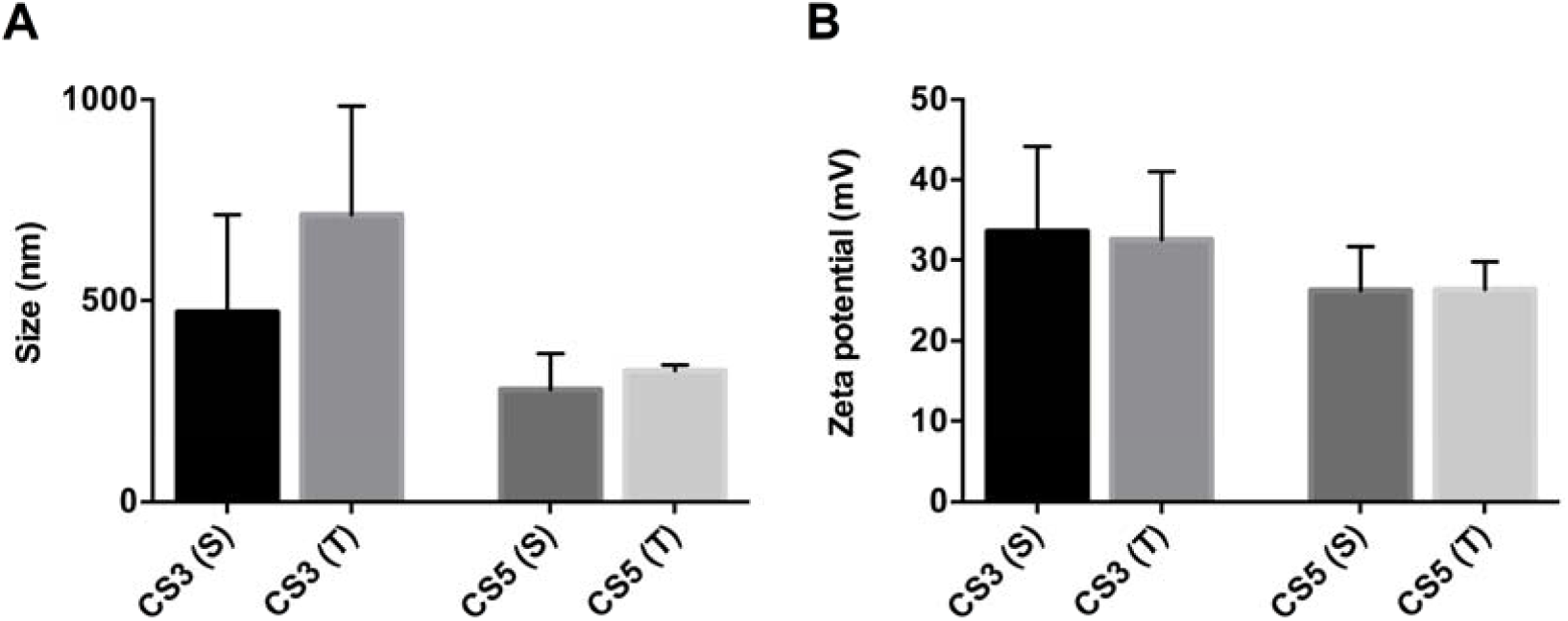
Comparison of physical properties: Size (graph A) and Zeta Potential (graph B) of CS3 (S) and CS3 (T), and CS5 (S) and CS5 (T) co-complexation polyplexes, with 100 µg of NLS-3 and NLS-5 peptides, respectively. Polydispersity was similar for all samples (below 0.4). Statistical differences between polyplexes with the same NLS peptide were calculated using Sidak’s multiple comparisons test. No differences were found between formulations. N=3. Error bars refer to standard deviation.

Despite a slight increase in polyplex mean size observed for (T) formulations compared to (S) formulations, no statistical differences were found for either NLS-3 or NLS-5 peptides. This increasing trend may be justified by the early interaction between DNA chains and NLS, which may have facilitated charge pairing during polyplex formation. The surface charge was also similar among formulations (Figure 2), indicating that the process of NLS addition does not significantly influence the physical properties of polyplexes.

To test for a possible effect in polyplex formation of the additional amount of positive charges brought by the addition of the NLS peptides in formulations CS-3 and CS-5, the formulations 15:1-CS3 and 15:1-CS5 (shown for 100 µg of NLS) were prepared considering the total amount of positively charged groups, including peptide and polymer, at a (NLS-3+CS):DNA or (NLS-5+CS):DNA (N:P ratio) of 15:1 and further characterized (Figure 3). Again, the addition of NLS was either totally made to the CS solution (T formulations) or split between this and DNA solutions (S formulations).

**Figure 3.**
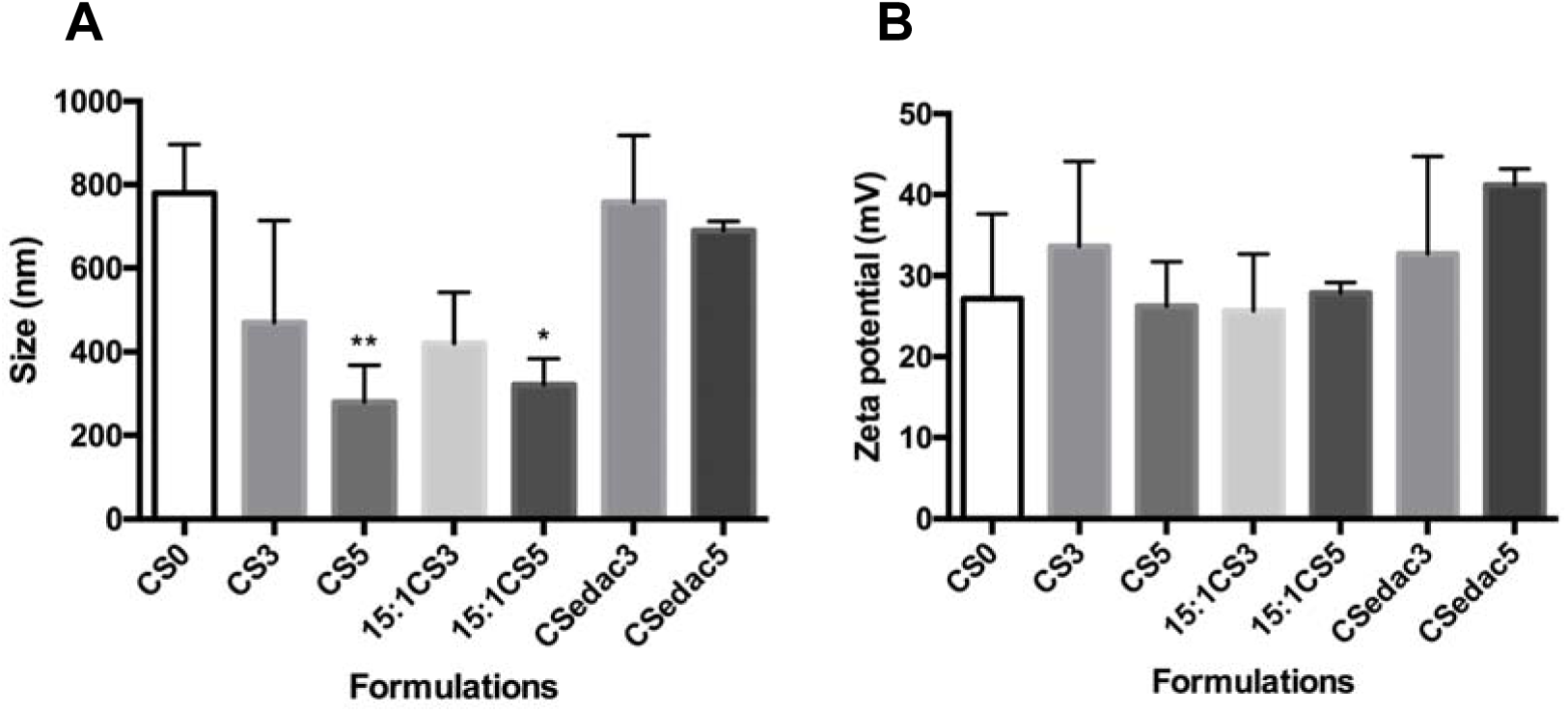
Comparison of physical properties: Size (graph A) and Zeta Potential (graph B) of polyplexes CS0, CS3, CS5, 15:1-CS3, 15:1-CS5, all with 100 µg of NLS (the graphs are representative for S and T co-complexation formulations) and covalently bond NLS formulations (CSedac3 and CSedac5). Polydispersity was below 0.4. Statistical differences of each condition compared to polyplexes without NLS peptides (CS0) were calculated using the Dunnett’s multiple comparisons test. N=3, error bars=SD; **p<0.01; *p<0.05.

Other than co-administration and co-complexation of NLS with polyplexes, we have also covalently linked the NLS to polyplexes. Several methods have been described in the literature to covalently bind NLS peptides to DNA:Ciolina *et al*. covalently associated NLS peptides to DNA by photoactivation, but these plasmid-NLS conjugates were not detected in the nucleus [44]; others have observed an increase on gene expression only when five NLS peptides were covalently coupled to DNA by diazo coupling through a PEG chain but not when the NLS peptides were directly coupled to DNA [45]; Zanta and colleagues covalently ligated one NLS-oligonucleotide to one or both ends of a linear DNA [46]. However, extensive chemical modification of DNA causes reduction or inhibition of gene expression [16, 22, 45, 47]. Therefore, to avoid the chemical modification of DNA, we have covalently linked NLS peptides to chitosan. This was achieved by first activating their terminal carboxylate with EDAC, followed by nucleophilic attack of CS amine groups. This strategy produced polyplex formulations CSedac3 and CSedac5, with physical properties as shown in figure 3 (size graph).

These polyplex formulations presented sizes larger than all other NLS formulations, and closer to those of the NLS free formulation, CS0. One possible explanation is the involvement of the carboxylate group in salt bridges with the ammonium and guanidium groups of lysine and arginine residues [42], thus less available for reaction. However, if it was the case, unreacted NLS would still be present and these formulations would resemble 15:1-CS3 and 15:1-CS5. Therefore, it is most likely that bonding occurred and that once linked to the CS chain, NLS are not able to play the same role as when they are added to the formulation in a free state. We had expected the surface charge to be of the same magnitude as other formulations with NLS, and that is confirmed (as observed in Figure 3, in the zeta potential graph).

The efficiency of cellular uptake and intracellular trafficking depends on the physical characteristics of polyplexes. A survey of the literature shows that larger particles get internalized less often than smaller particles but have a higher rate of gene release into the cytosol due to their prolonged residence time. This prolonged residence time is an indication that larger particles most likely avoid rapid lysosomal degradation. Particles with sizes from 200 nm to 1 µm are internalized mainly by caveolae-mediated endocytosis [48], and the motility of caveolae is relatively low, depending on the network of actin filaments and microtubules [49]. Considering all our polyplexes have a size distribution ranging from 250 to 750 nm, these results suggest the predominant internalization pathway for our polyplexes to be caveolae-mediated endocytosis. The surface charge also affects cellular uptake of polyplexes, and according to the literature, positively charged polyplexes interact efficiently with the cell membrane [36]. All formulations in our study are positively charged polyplexes, thus favoring internalization.

### 3.2. DNA is efficiently complexed in the presence of NLS

DNA complexation in polyplex formulations was evaluated by agarose gel electrophoresis. DNA complexation was effectively achieved for all formulations, confirmed by the absence of free DNA migration in the gel (Figure 4), regardless of the NLS peptide.

**Figure 4.**
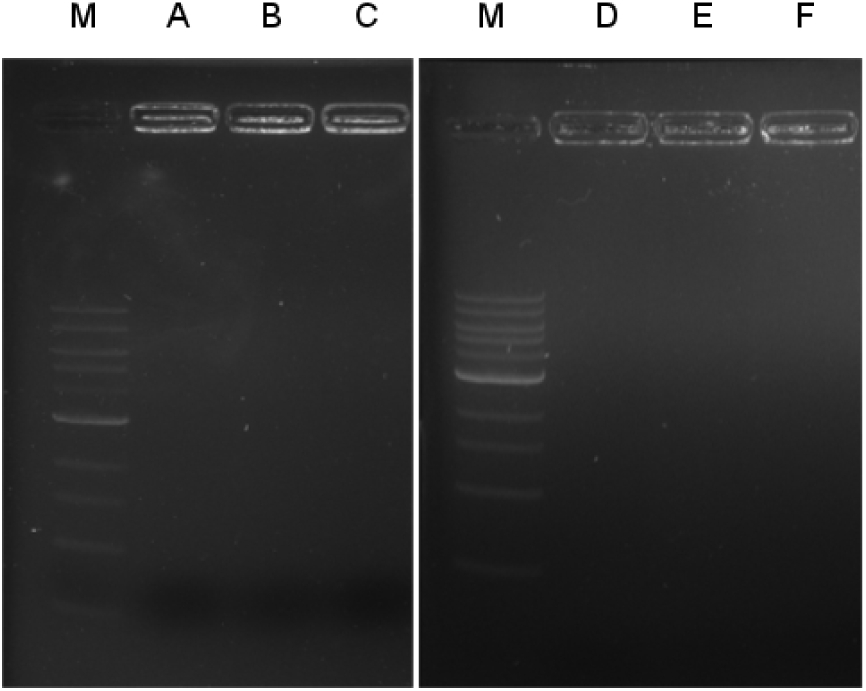
Representative images of the evaluation of DNA complexation by polyplexes by agarose gel electrophoresis and visualized by GreenSafe Premium. Polyplexes display efficient DNA complexation, evidenced by the absence of free DNA migration on the agarose gel. M – DNA Marker; A – CS3 (T) 100 µg; B – CS5 (T) 100 µg; C – 15:1CS3 (T); D – 15:1CS3 (S); E – 15:1CS5 (T); F – 15:1CS5 (S).

### 3.3. NLS are cytocompatible

The cytotoxicity of NLS peptides was evaluated at 24h and 72h using HEK293T cells and the MTT assay, as depicted in figure 5. No cytotoxicity was observed for either NLS-3 and NLS-5 peptides regardless of the amount, since cell viability was above 90% for all tested concentrations.

**Figure 5.**
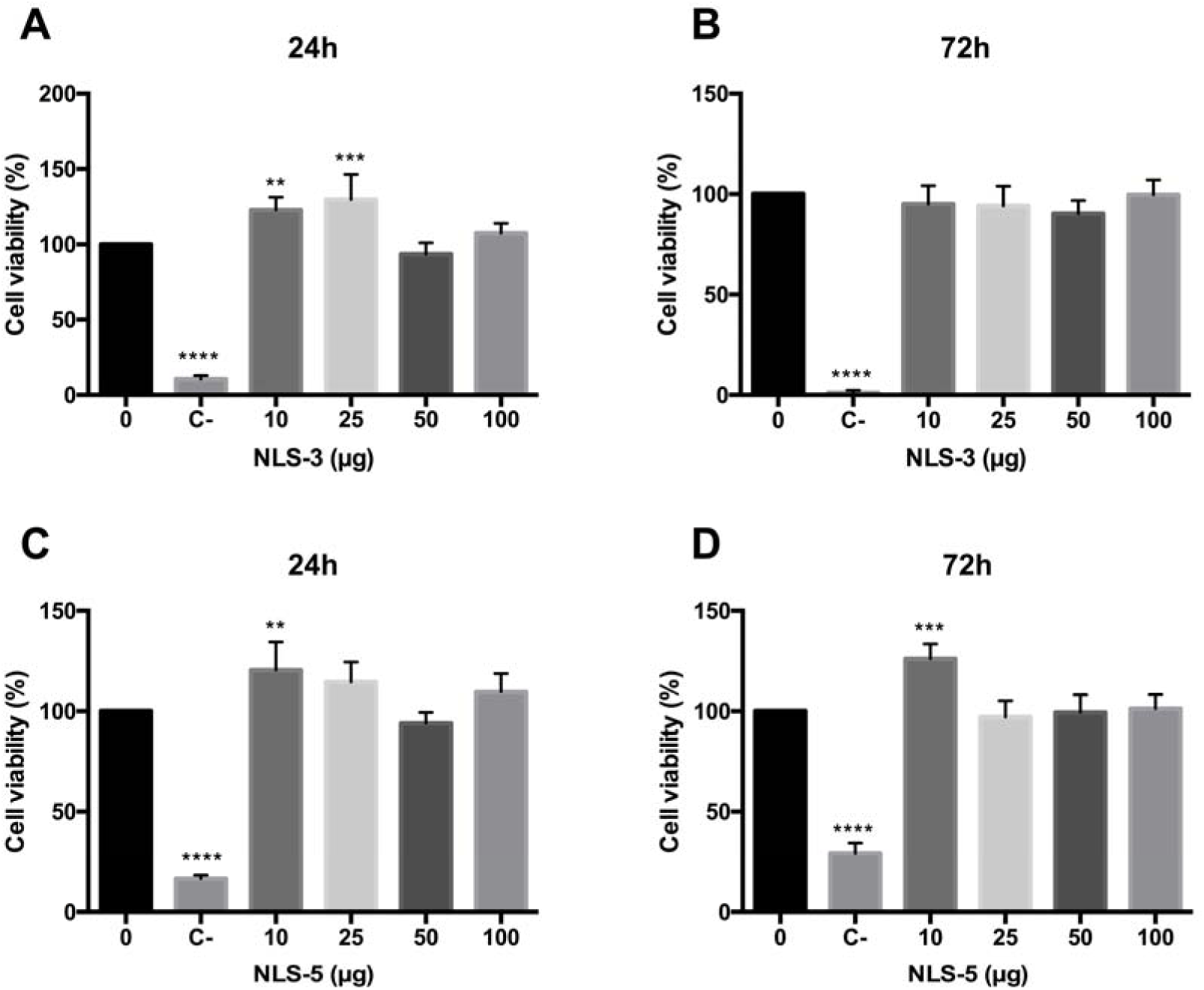
Cell viability (%) after 24h and 72h of incubation with different amounts of NLS-3 (graphs A, B) and NLS-5 (graphs C, D) peptides. Untreated cells were used as positive viability control (0 µg) and cells incubated with latex extracts as negative viability control (C-). Statistical differences, compared to positive control (0 µg), were calculated using Dunnett’s multiple comparisons test (****p<0.0001; ***p<0.001; **p<0.01). N=3, error bars=SD.

### 3.4. Improvement in transfection efficiency is NLS and method dependent

The transfection efficiency of all formulations was evaluated in HEK293T cells, which is a commonly used cell line for transfection studies. The gene transfer efficiency of the different NLS incorporation strategies was evaluated by GFP expression, visualized by fluorescence microscopy after 48h and 72h (supplementary data), and quantitatively analyzed by flow cytometry at 72h (Figures 6 to 9).

**Figure 6.**
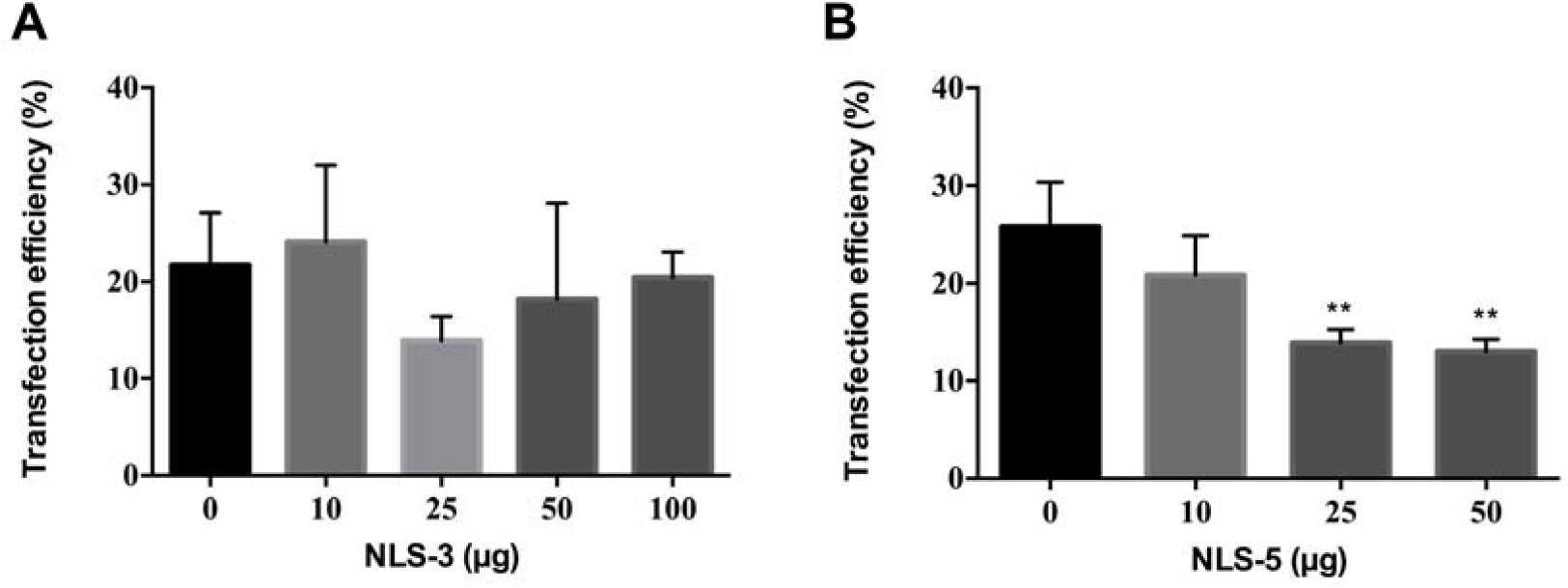
Transfection efficiency expressed as percentage of GFP-positive cells after co-administration of polyplexes (CS) with NLS-3 (graph A) or NLS-5 (graph B). Transfection was performed with 1 µg of DNA for all groups and analyzed 72h after transfection. N=3, bars correspond to SD. Statistical significance was tested by one-way ANOVA with Dunnett’s post-test compared against polyplexes without NLS peptides (0 µg) (**p<0.01).

#### 3.4.1. Co-administration of polyplexes and NLS does not improve transfection efficiency

To evaluate the transfection efficiency of NLS co-administered with polyplexes the formulation CS0 and NLS solution were simultaneously added at the time of transfection. For NLS-3 peptide amounts between 10 and 150 µg were co-administrated and for NLS-5 the range varied between 10 and 50 µg, since no improvement was observed with higher concentrations of peptides in preliminary studies [50].

The *in vitro* transfection ability of CS polyplexes without NLS peptides (Figure 6) was relatively low, reaching 25% transfection efficiency. Similar results were achieved in a study with the same preparation method of polyplexes in HEK293 cells, where a low transfection efficiency of chitosan/DNA complexes was obtained [30].

Contrary to what has been described in the literature for other NLS, no increase in transfection efficiency was observed when either peptide was co-administered with CS0 polyplexes, regardless of the amounts of NLS (Figure 6: conditions 10, 25, 50, and 100 µg), when compared with polyplexes without NLS peptides (Figure 6, condition 0 µg). Moreover, polyplexes with IGFBP-5 peptides registered a significant decrease in transfection when compared with polyplexes without NLS peptides. In spite of polyplexes having size and surface charge appropriate for gene delivery, and the cationic NLS peptides containing lysine and arginine and thus being capable of binding to DNA through electrostatic interactions [19], in our study co-administration has not improved transfection efficiency.

#### 3.4.2. Co-complexation of polyplexes and NLS improves transfection efficiency

NLSs can be coupled to DNA or to vectors to improve gene delivery but it is not yet clear what is the best method to incorporate the NLS peptides [51]. Yoo *et al* attached psoralen-NLS conjugates non-covalently to DNA/PEI complexes and observed an increase in transfection efficiency in COS-1 cells when compared to a mutant NLS or DNA/PEI complexes without NLS peptides [47]. Other authors incorporated NLS peptides directly into DNA or polymer, without covalent conjugation. The polyplexes with increasing ratio of co-complexed NLS peptides increased the transfection efficiency in HeLa cells, in a NLS-dose dependent manner [22].

We have therefore prepared polyplexes by co-complexation of chitosan, DNA, and increasing amounts of NLS-3 and NLS-5 peptides. To test if the NLS incorporation strategy influenced the transfection efficiency, we have prepared polyplexes with total NLS added to chitosan [CS3 (T), CS5 (T)] and NLS equally divided between chitosan and DNA solutions [CS3 (S), CS5 (S)]. GFP expression was quantified as shown in Figure 7, which is representative of both strategies (total and split).

**Figure 7.**
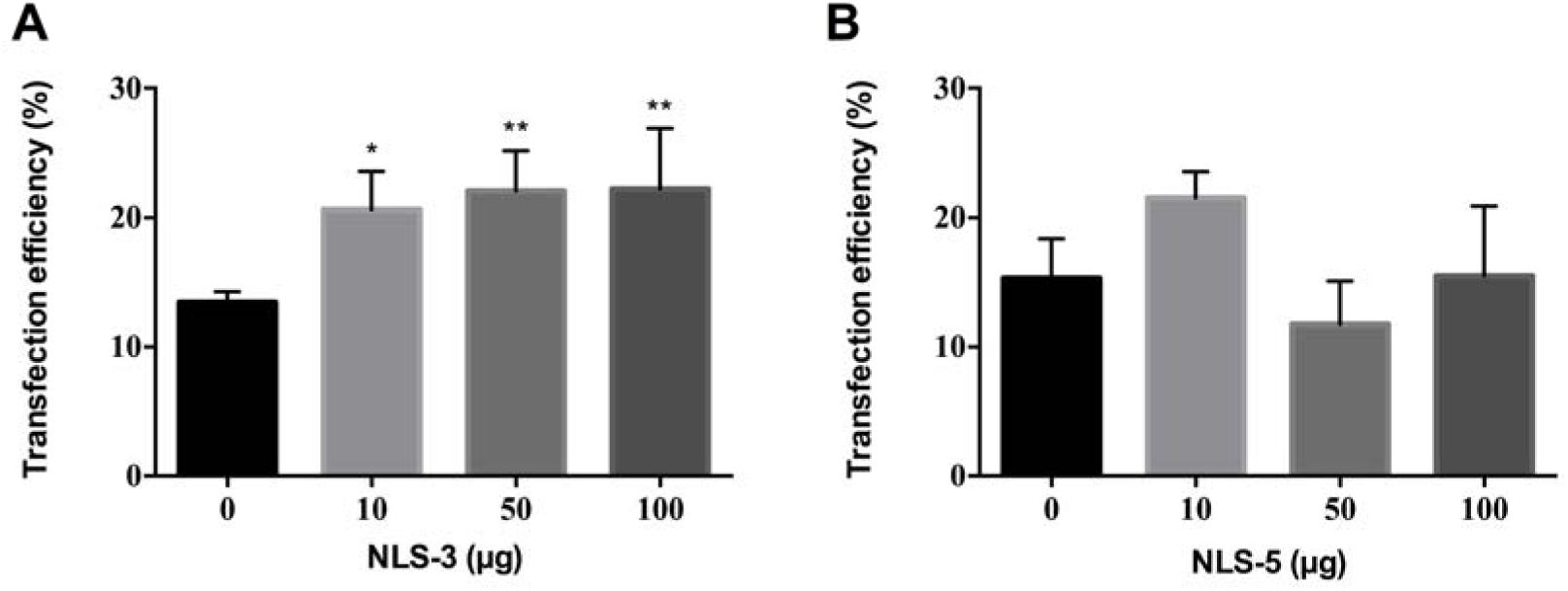
Transfection efficiency expressed as percentage of GFP-positive cell for CS3 (S) (graph A) and CS5 (S) (graph B) co-complexation polyplexes. Transfection was performed with 1 µg of DNA for all groups and analyzed 72h after transfection. N=3, bars correspond to SD. Statistical differences were calculated using Dunnett’s multiple comparisons test compared with polyplexes without NLS peptides (0 µg) (**p<0.01; *p<0.05).

A significant increase in transfection efficiency was observed with polyplexes co-complexed with NLS-3 peptides when compared to polyplexes without NLS (CS, 0 µg). Contrary to what was expected, this trend was not observed in polyplexes co-complexed with NLS-5 peptides. This was observed for both incorporation strategies (total and split) and although this had an effect on the size of polyplexes (Figure 2), it does not appear to affect their efficiency, as reported for other systems [35].

Previous studies reported that the affinity or accessibility to importin subunits differ between IGFBP-3 and IGFBP-5, although it is not clear how [26]. This could explain the difference in the results of Figure 7. When NLS-5 peptides are co-complexed with CS polyplexes, the accessibility of the peptides to their nuclear receptors (i.e. importins) might be hampered and DNA nuclear delivery is lower, hence no improvement in transfection efficiency is observed.

Although the addition of NLS peptides did not significantly alter the size or surface charge of polyplexes, we have nonetheless tested if formulations accounting for NLS charge addition (15:1-CS3 and 15:1-CS5) had an impact in transfection efficiency (Figure 8). We have chosen a single amount of NLS (100 µg) to evaluate potential differences between formulations.

**Figure 8.**
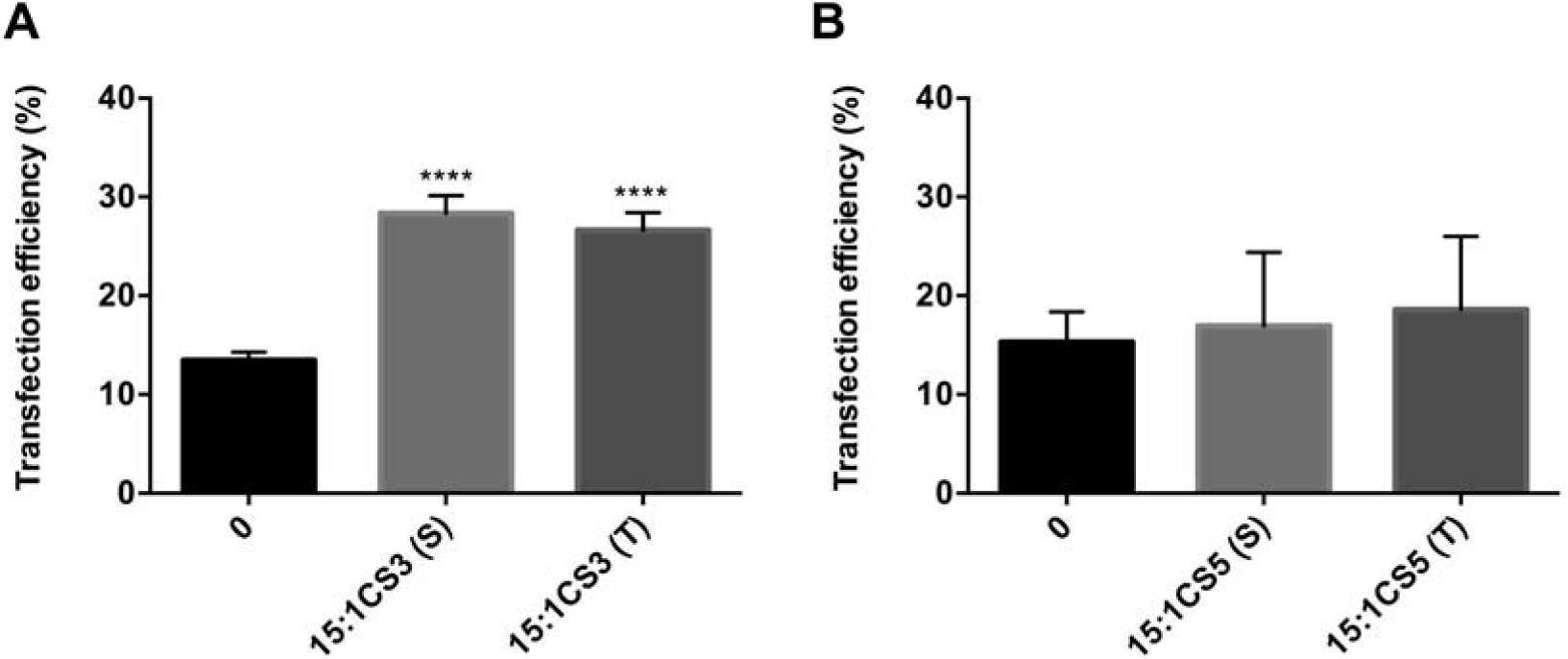
Transfection efficiency expressed as percentage of GFP-positive cells by 15:1-CS3 (S) and (T) (graph A) and 15:1-CS5 (S) and (T) (graph B) co-complexation polyplexes with 100 µg of NLS peptides. Transfection was performed with 1 µg of DNA for all groups and analyzed 72h after transfection. N=3, bars correspond to SD. Statistical differences compared with CS polyplexes (condition 0 in both graphs) were calculated using Dunnett’s multiple comparisons test (****p<0.0001).

The effect of N:P ratio of polyplexes has been widely investigated [19, 20, 22, 30, 35] and described as influencing polyplex formation (size and surface charge) and transfection efficiency [36]. In this work, the amount of chitosan was adjusted to include peptides in the N:P ratio and therefore to maintain the number of amine groups available for interaction with the DNA. Hence, the statistically significant 2-fold increase in transfection efficiency observed for 15:1CS3 polyplexes, (S) and (T), when compared with polyplexes without NLS peptides (condition 0, figure 8, graph A) is attributed to the presence of NLS. For polyplexes with NLS-5 peptides, no increase in transfection efficiency was observed when compared to CS complexes (figure 8, graph B).

#### 3.4.3. Covalent ligation of NLS to polyplexes improves transfection efficiency

The third method chosen for the incorporation of NLS peptides was covalent ligation. Several methods for covalently linking molecules to DNA have been developed [44-46, 52] but the chemical modification of DNA might cause a decrease in its transcription [16, 47]. In this study, we have therefore chosen to covalently link NLS peptides to the polymer. This was performed via amide bond formation between the carboxylic acid moieties of the NLS peptides and the amine groups of chitosan, which was mediated by a carbodiimide (EDAC). Resulting polyplexes were also tested for their transfection efficiency, as shown in Figure 9.

**Figure 9.**
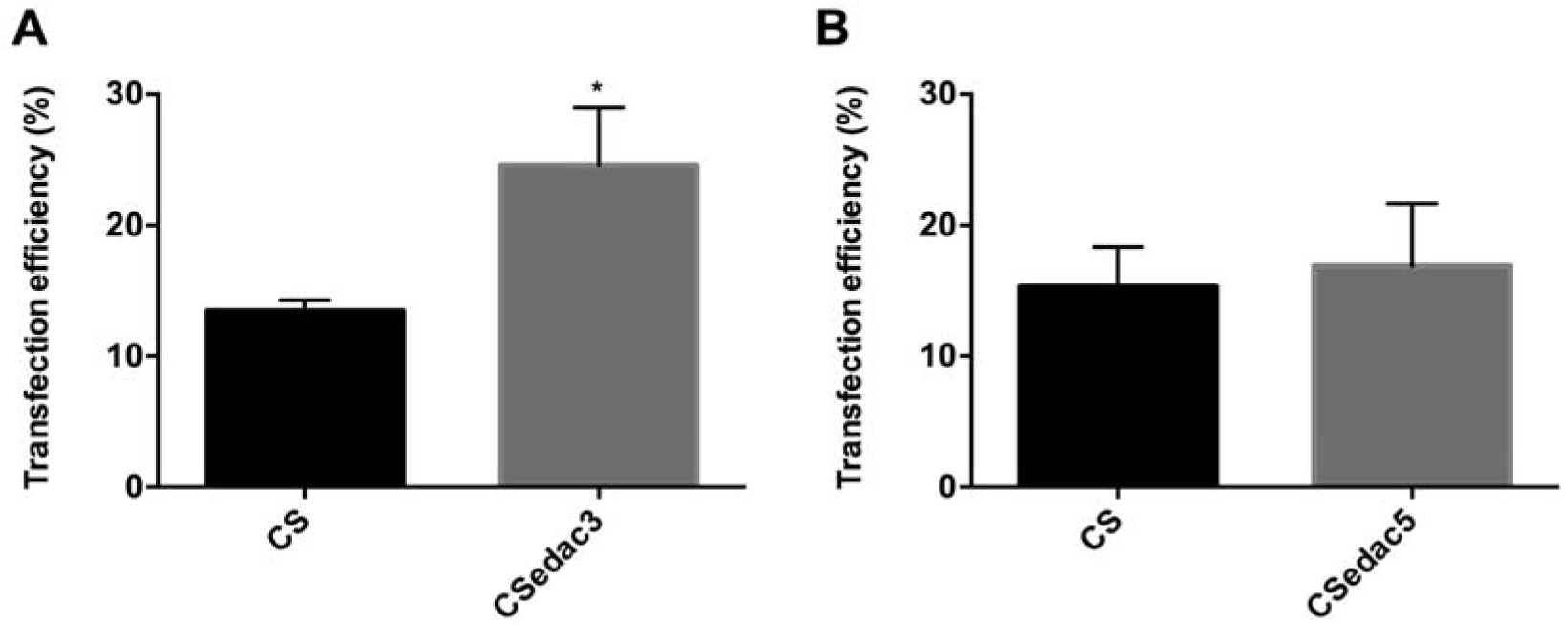
Transfection efficiency represented by percentage of GFP-positive cells of CSedac3 (graph A) and CSedac5 (graph B) polyplexes covalently linked with 100 µg of NLS peptides. Transfection was performed with 1 µg of DNA for all groups and analyzed 72h after transfection. N=3, bars correspond to SD. Statistical differences compared with CS polyplexes were calculated using an Unpaired t test with Welch’s correction (*p<0.05).

In figure 9 a 2-fold enhancement in the transfection efficiency of CSedac3 polyplexes is observed when compared to polyplexes without NLS peptides. Again, no increase in transfection efficiency was observed for CSedac5 polyplexes compared to polyplexes without NLS peptides. These results indicate that transfection efficiency is improved by polyplexes with IGFBP-3 peptides but not with polyplexes with IGFBP-5 peptides, suggesting, as hypothesized above, that recognition of IGFBP-5 peptides by importins may be hampered by their entanglement during polyplex preparation. This is supported by the results for polyplex size, which are smaller for formulations with IGFBP-5 peptides, a reflection of a higher degree of entanglement.

When compared with the other two methodologies (co-administration and co-complexation) CSedac3 and CSedac5 polyplexes yielded similar transfection efficiencies to co-complexation.

## Conclusions

The goal of this study was to determine if the transfection efficiency of chitosan-based non-viral gene delivery systems can be improved through the incorporation of NLS peptides. We have tested three approaches to incorporate the IGFBP-derived NLS peptides into polyplexes: co-administration at the time of transfection, co-complexation during polyplex production and covalent ligation to the polymer component prior to polyplex preparation. The characterization of the resulting formulations has shown that sodium sulfate has a role in polymer-DNA entanglement, since its addition yielded polyplexes with smaller size and surface charge.

We have also observed that the addition of increasing amounts of NLS peptides to polyplexes influenced their physical properties, such as size and surface charge, which were NLS dependent. However, regardless of the incorporation technique, the presence of sodium sulfate, N:P ratio, or addition of NLS peptides, all polyplex formulations were positively charged, capable of effective DNA complexation and with a size suitable for their use for gene delivery. The NLS peptides were not cytotoxic to HEK293T cells and lastly, *in vitro* transfection assays showed that transfection efficiency of polyplexes was dependent on the NLS peptide and incorporation strategy. Co-administration of the polyplexes and NLS-5 showed no improvement in transfection efficiency. On the other hand, formulations of co-complexed or covalently attached NLS-3 peptides had a 2-fold increase in transfection efficiency, most likely due to higher affinity with or accessibility to importin subunits between NLS-3 and NLS-5 peptides.

To the best of our knowledge, this is the first study where NLS peptides derived from human molecules are associated to chitosan-based non-viral gene vectors. Overall the results shown that polyplexes co-complexed with NLS-3 peptides are indeed good candidates for non-viral gene delivery systems. Further studies will include their evaluation using a relevant gene, cellular model and further on in an animal model.

## Acknowledgments

The authors acknowledge Dr. Jean Bennett (University of Pennsylvania, USA) for kindly providing the GFP plasmid and Dr. Lyn Schedlich and Sue Firth for kindly providing the NLS plasmids. The authors also acknowledge the financial support of Fundação para a Ciência e Tecnologia (PTDC/SAU-BEB/098475/2008 to Gabriela A. Silva, PD/BD/52424/2013 individual fellowship to Diogo Bitoque, SFRH/BD/70318/2010 individual fellowship to Ana V. Oliveira, SFRH/BD/76873/2011 individual fellowship to Sofia Calado, SFRH/BPD/78404/2011 individual fellowship to Sónia Simão, IBB/LA under the project PEst-OE/EQB/LA0023/2013, and PEst-OE_QUI_UI4023_2011) and the Marie Curie Reintegration Grant (PIRG-GA-2009-249314 to Gabriela A. Silva) under the FP7 program. iNOVA4Health - UID/Multi/04462/2013, a program financially supported by Fundação para a Ciência e Tecnologia / Ministério da Educação e Ciência, through national funds and co-funded by FEDER under the PT2020 Partnership Agreement is also acknowledged.

## Contribuitions

Diogo B. Bitoque* wrote the manuscript, Joana Morais* and Ana V. Oliveira* performed the experiments, Raquel L. Sequeira reviewed the manuscript, Sofia M. Calado and Tiago M. Fortunato performed the cloning experiments, Sónia Simão revised the manuscript, Ana M. Rosa da Costa designed the chemistry experiments and revised the manuscript and Gabriela A. Silva designed the project, funded and revised the manuscript.

