## Supplemental Data for "Human-derived NLS enhance the gene transfer efficiency of chitosan"

Supplementary data:

**B**
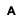

**A**

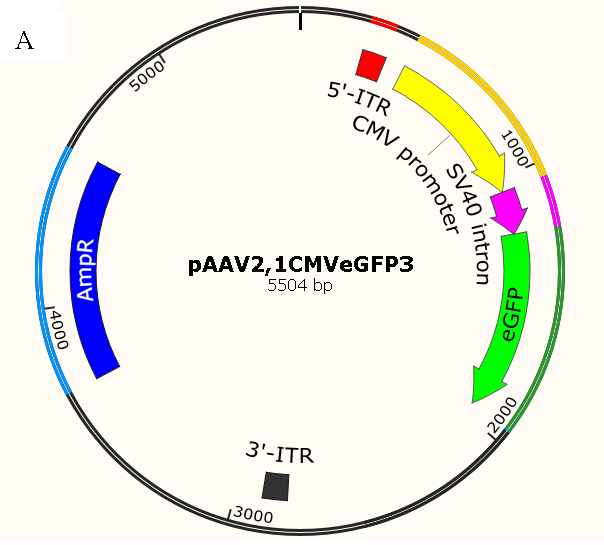

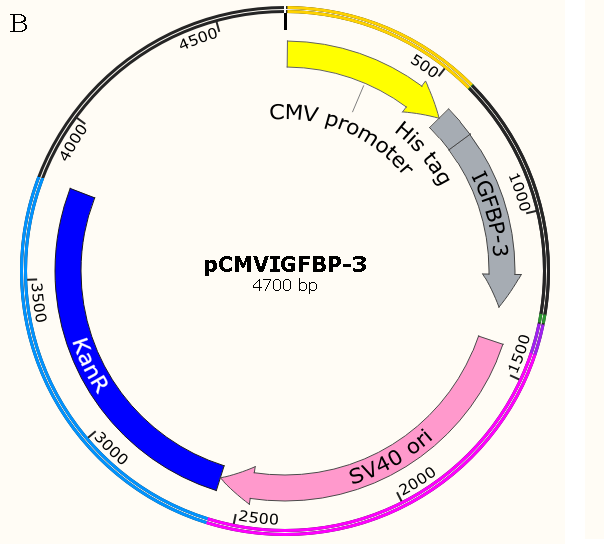

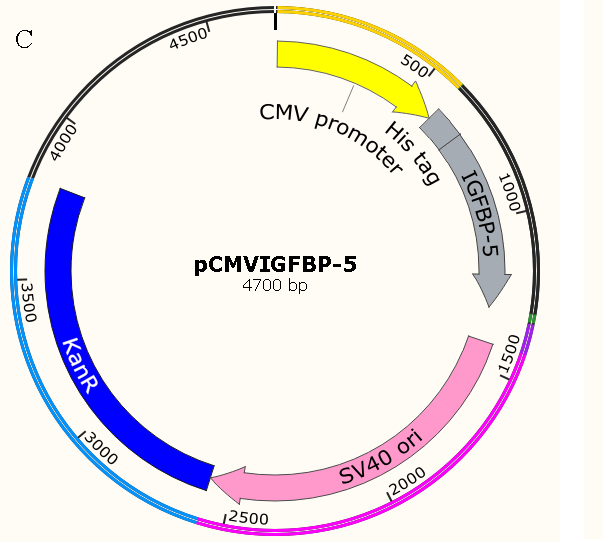

**C**

Figure S1 - Schematic representation of the structure of plasmids. (A) pAAV2.1CMVeGFP3 used for expression of GFP (B) pCMVIGFBP-3 used to encode IGFBP-3 peptide and (C) pCMVIGFBP-5 used to encode IGFBP-5 peptide. AmpR and KanR are genes for resistance to ampicilin and kanamycin, respectively.

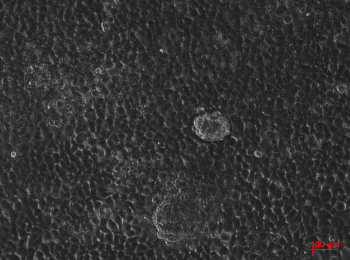

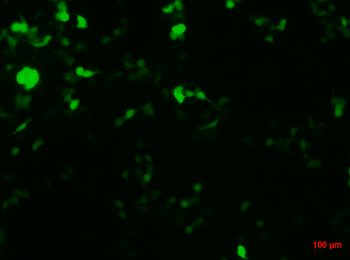

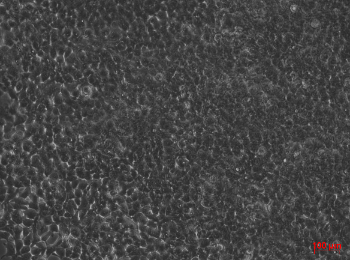

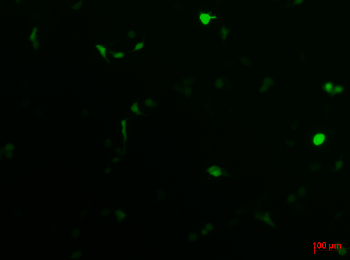

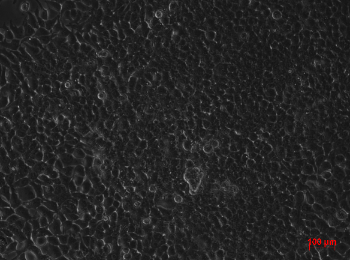

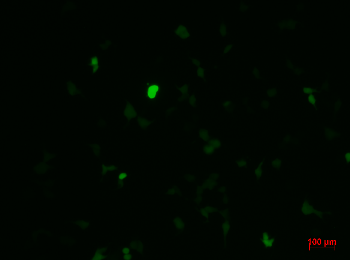

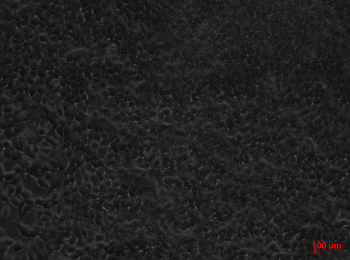

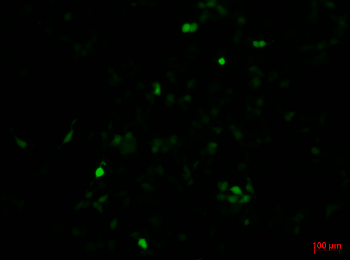

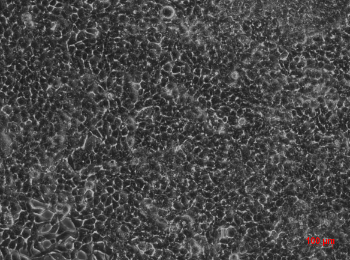

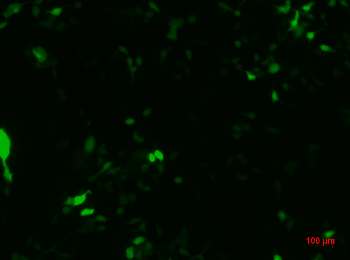

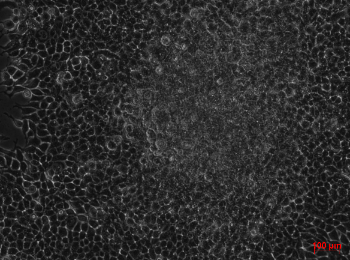

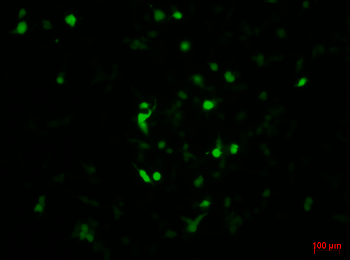

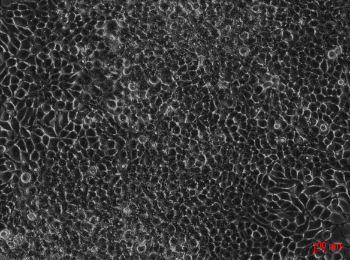

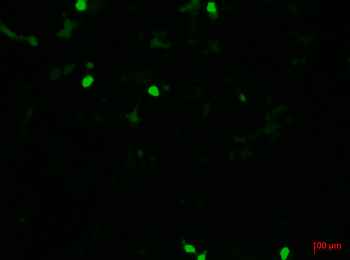

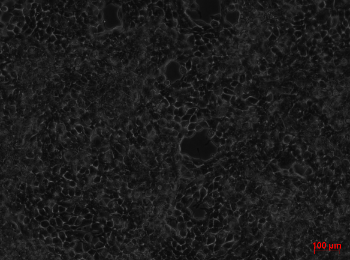

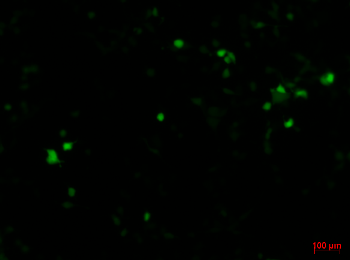

**Bright field**

**1**

**1**

**1**

**1**

**1**

**1**

**1**

**Fluorescence**

**2**

**2**

**2**

**2**

**2**

**2**

**2**

**Bright field**

**1**

**1**

**1**

**1**

**1**

**1**

**1**

**Fluorescence**

**2**

**2**

**2**

**2**

**2**

**2**

**2**

**25 µg**

**A**

**A**

**A**

**A**

**A**

**A**

**A**

**50 µg**

**A**

**A**

**A**

**A**

**A**

**A**

**A**

**100 µg**

**A**

**A**

**A**

**A**

**A**

**A**

**A**

**10 µg**

**A**

**A**

**A**

**A**

**A**

**A**

**A**

**A**

**B**

Figure S2 - Representative images of fluorescence microscopy of transfected cells by CSNa_2_SO_4_ polyplexes with several concentrations of IGFBP-3, after 48h (A) and 72h (B). Amplification of 100X and scale bar represents 100 µm

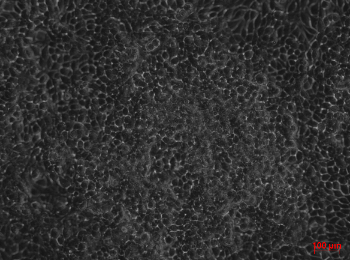

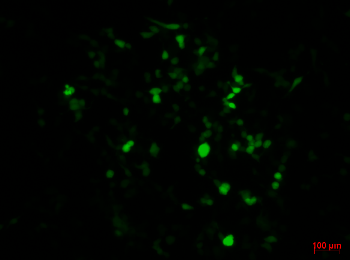

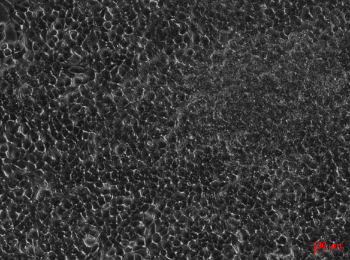

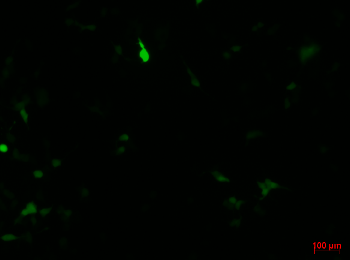

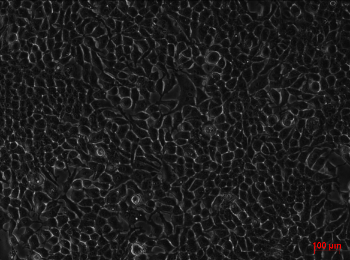

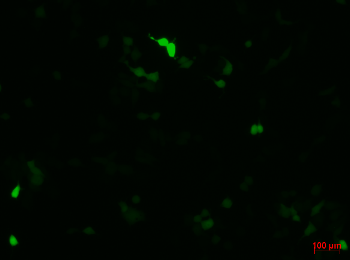

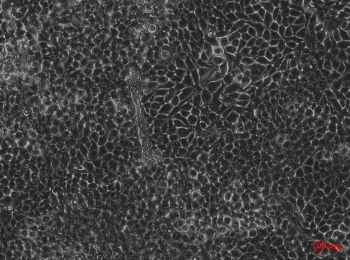

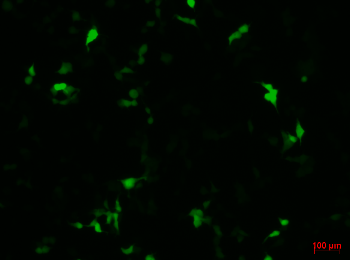

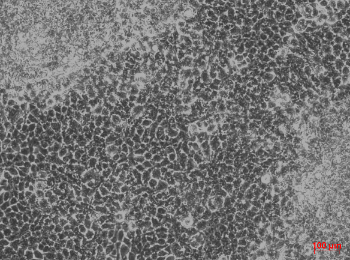

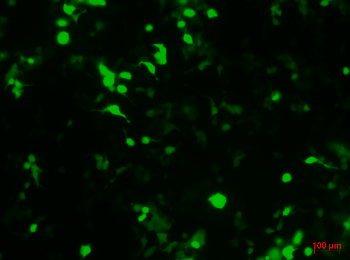

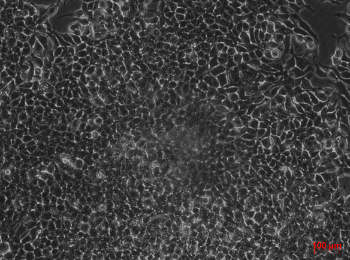

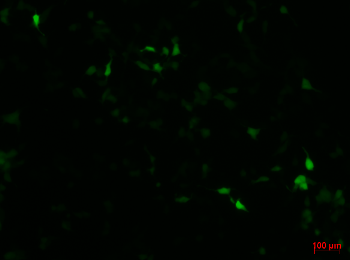

**10 µg**

**A**

**A**

**A**

**A**

**A**

**A**

**A**

**25 µg**

**A**

**A**

**A**

**A**

**A**

**A**

**A**

**50 µg**

**A**

**A**

**A**

**A**

**A**

**A**

**A**

**Bright field**

**1**

**1**

**1**

**1**

**1**

**1**

**1**

**Fluorescence**

**2**

**2**

**2**

**2**

**2**

**2**

**2**

**Bright field**

**1**

**1**

**1**

**1**

**1**

**1**

**1**

**Fluorescence**

**2**

**2**

**2**

**2**

**2**

**2**

**2**

**A**

**B**

Figure S3 - Representative images of fluorescence microscopy of transfected cells by CSNa_2_SO_4_5 polyplexes with several concentrations of IGFBP-5 peptides, after 48h (A) and 72h (B). Amplification of 100X and scale bar represents 100 µm.

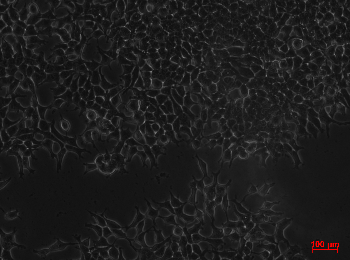

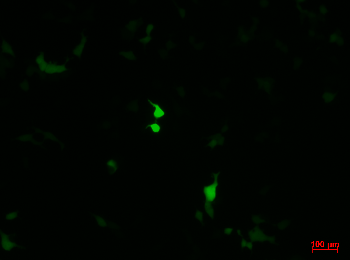

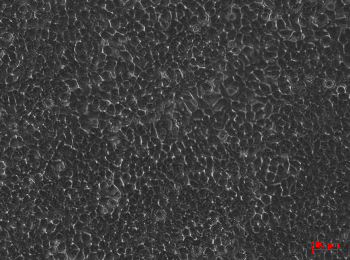

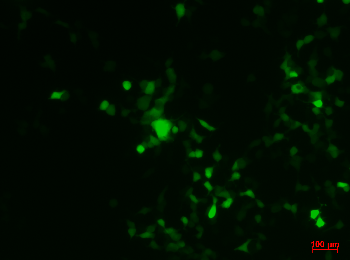

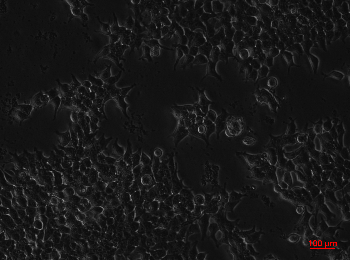

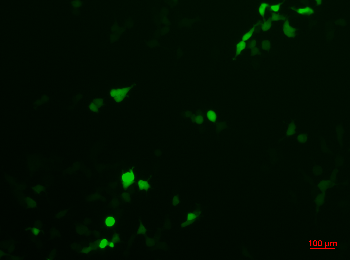

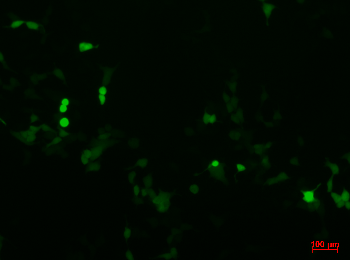

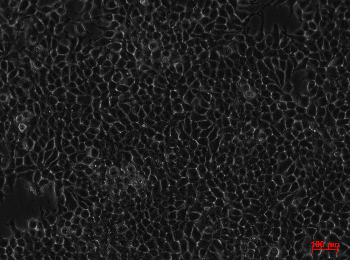

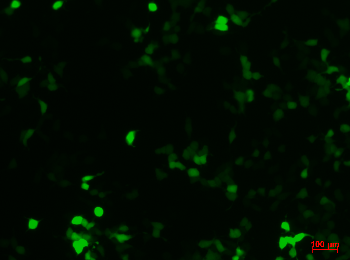

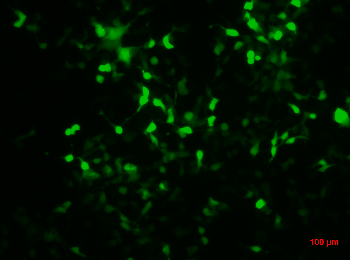

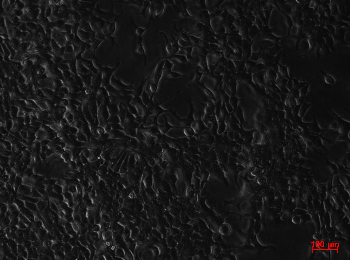

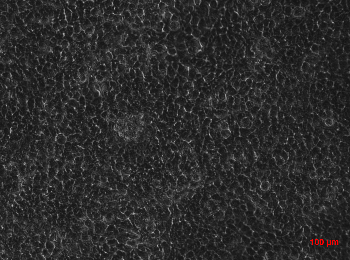

**10 µg**

**A**

**A**

**A**

**A**

**A**

**A**

**A**

**50 µg**

**A**

**A**

**A**

**A**

**A**

**A**

**A**

**100 µg**

**A**

**A**

**A**

**A**

**A**

**A**

**A**

**Bright field**

**1**

**1**

**1**

**1**

**1**

**1**

**1**

**Fluorescence**

**2**

**2**

**2**

**2**

**2**

**2**

**2**

**Bright field**

**1**

**1**

**1**

**1**

**1**

**1**

**1**

**Fluorescence**

**2**

**2**

**2**

**2**

**2**

**2**

**2**

**A**

**B**

Figure S4 - Representative images of fluorescence microscopy of transfected cells by CS3 (S) polyplexes with several concentrations of IGFBP-3, after 48h (A) and 72h (B). Amplification of 100X and scale bar represents 100 µm.

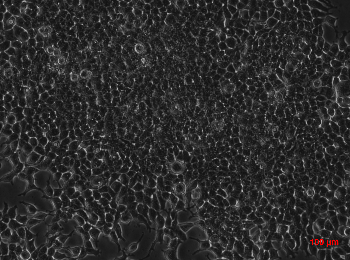

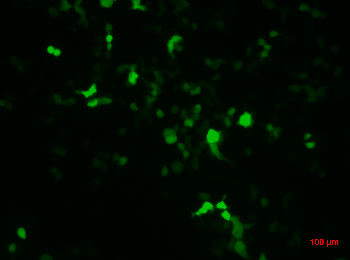

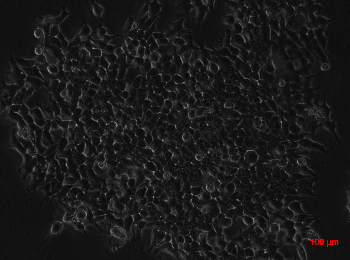

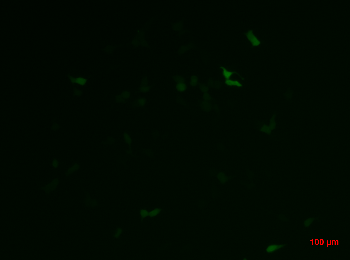

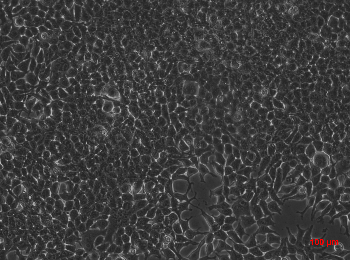

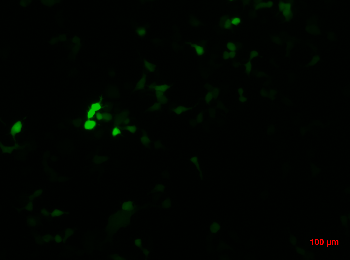

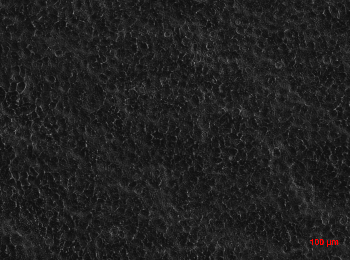

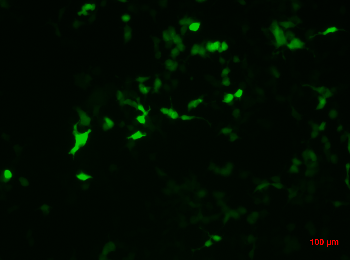

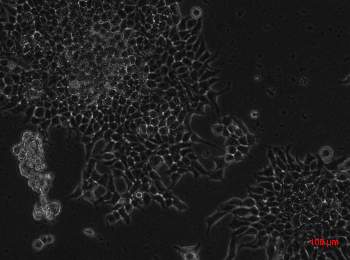

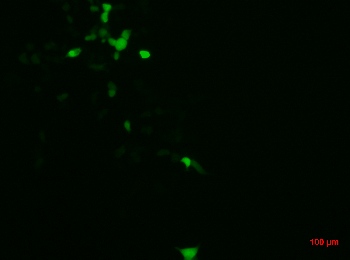

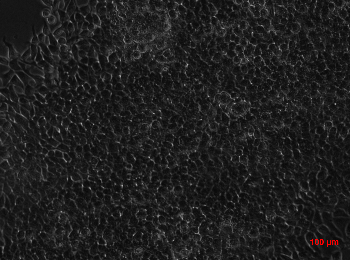

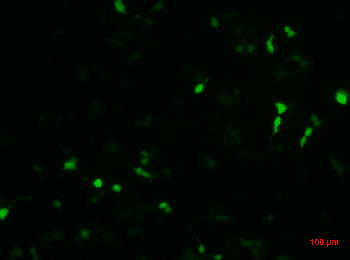

**10 µg**

**A**

**A**

**A**

**A**

**A**

**A**

**A**

**50 µg**

**A**

**A**

**A**

**A**

**A**

**A**

**A**

**100 µg**

**A**

**A**

**A**

**A**

**A**

**A**

**A**

**Bright field**

**1**

**1**

**1**

**1**

**1**

**1**

**1**

**Fluorescence**

**2**

**2**

**2**

**2**

**2**

**2**

**2**

**Bright field**

**1**

**1**

**1**

**1**

**1**

**1**

**1**

**Fluorescence**

**2**

**2**

**2**

**2**

**2**

**2**

**2**

**A**

**B**

Figure S5 - Representative images of fluorescence microscopy of transfected cells by CS5 (S) polyplexes with several concentrations of IGFBP-5, after 48h (A) and 72h (B). Amplification of 100X and scale bar represents 100 µm.

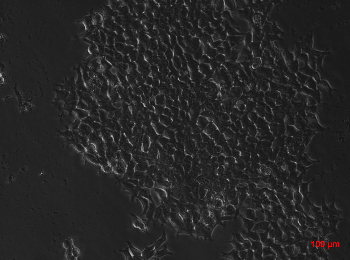

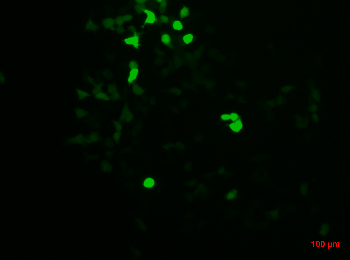

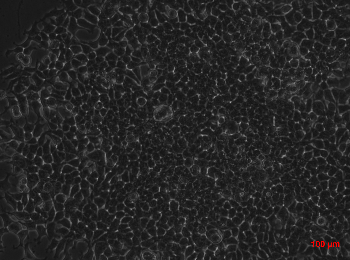

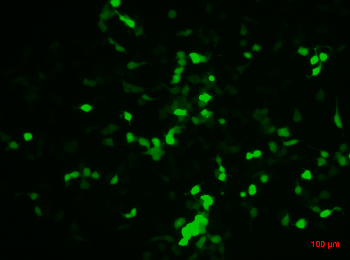

**CS5 (T)**

**B**

**B**

**B**

**B**

**B**

**B**

**B**

**CS3 (T)**

**A**

**A**

**A**

**A**

**A**

**A**

**A**

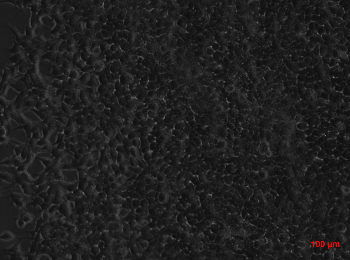

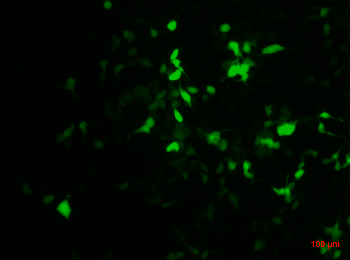

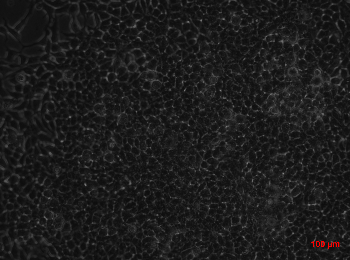

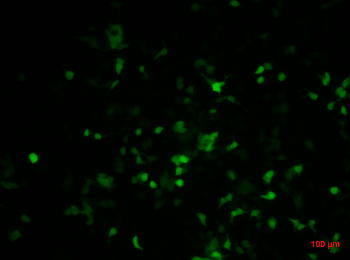

**Bright field**

**1**

**1**

**1**

**1**

**1**

**1**

**1**

**Fluorescence**

**2**

**2**

**2**

**2**

**2**

**2**

**2**

**Bright field**

**1**

**1**

**1**

**1**

**1**

**1**

**1**

**Fluorescence**

**2**

**2**

**2**

**2**

**2**

**2**

**2**

**A**

**B**

Figure S6 - Representative images of fluorescence microscopy of transfected cells by CS3 and CS5, (T), polyplexes with 100 µg of IGFBP-3 or IGFBP-5, respectively. Cells were visualized after 48h (A) and 72h (B). Amplification of 100X and scale bar represents 100 µm.

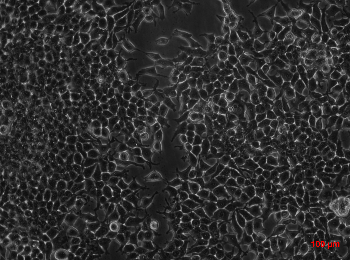

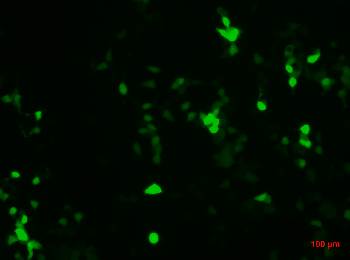

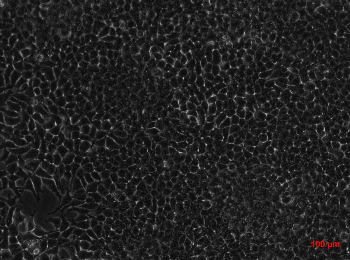

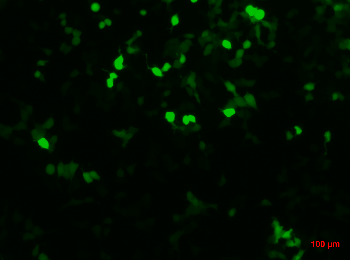

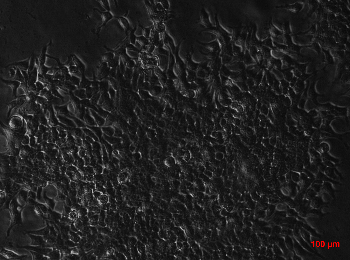

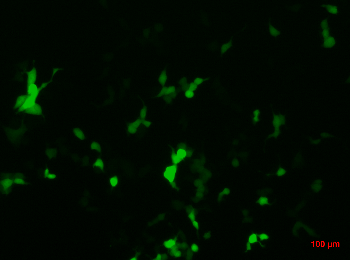

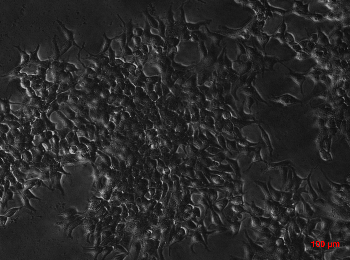

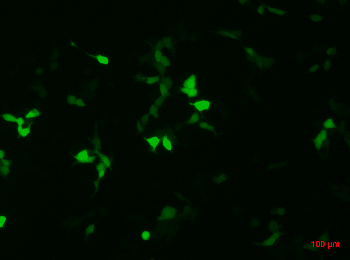

**15:1CS3**

**(T)**

**15:1CS3**

**(S)**

**Bright field**

**1**

**1**

**1**

**1**

**1**

**1**

**1**

**Fluorescence**

**2**

**2**

**2**

**2**

**2**

**2**

**2**

**Bright field**

**1**

**1**

**1**

**1**

**1**

**1**

**1**

**Fluorescence**

**2**

**2**

**2**

**2**

**2**

**2**

**2**

**A**

**B**

Figure S7 - Representative images of fluorescence microscopy of transfected cells by 15:1CS3 (S) and 15:1CS3 (T) polyplexes with 100 µg of IGFBP-3, after 48h (A) and 72h (B). Amplification of 100X and scale bar represents 100 µm.

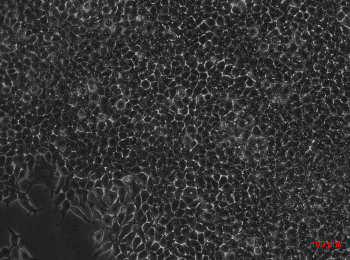

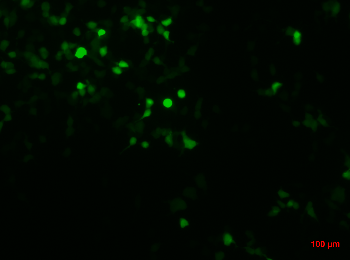

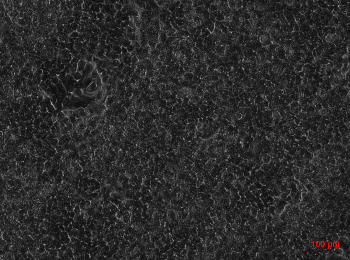

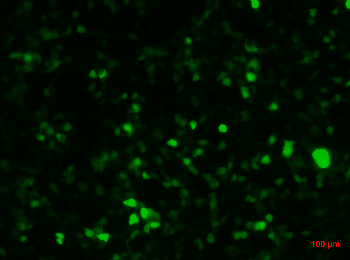

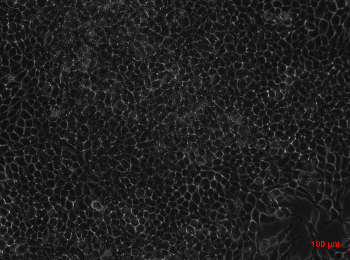

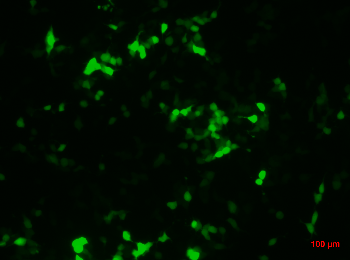

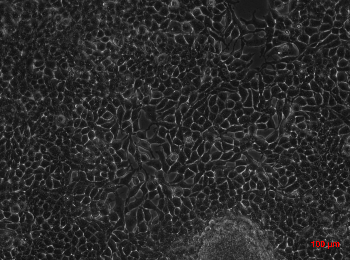

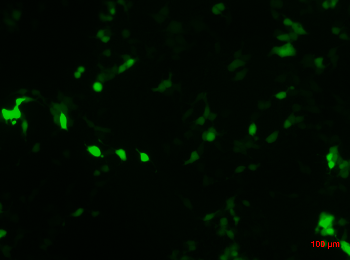

**15:1CS5**

**(T)**

**15:1CS5**

**(S)**

**Bright field**

**1**

**1**

**1**

**1**

**1**

**1**

**1**

**Fluorescence**

**2**

**2**

**2**

**2**

**2**

**2**

**2**

**Bright field**

**1**

**1**

**1**

**1**

**1**

**1**

**1**

**Fluorescence**

**2**

**2**

**2**

**2**

**2**

**2**

**2**

**A**

**B**

Figure S8 - Representative images of fluorescence microscopy of transfected cells by 15:1CS5 (S) and 15:1CS5 (T) polyplexes with 100 µg of IGFBP-5, after 48h (A) and 72h (B). Amplification of 100X and scale bar represents 100 µm.

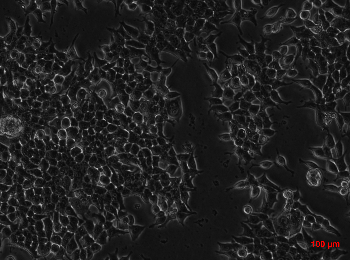

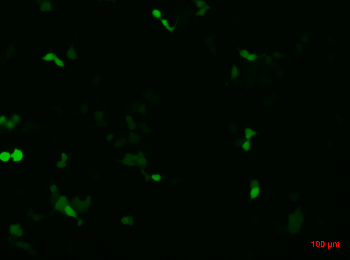

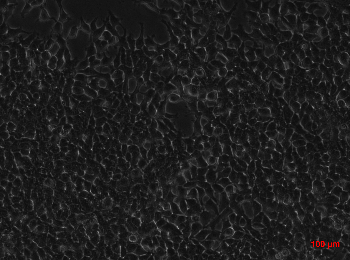

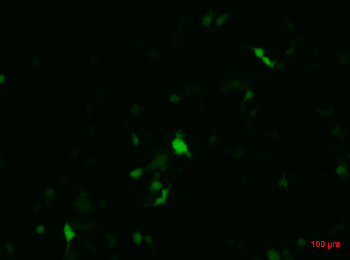

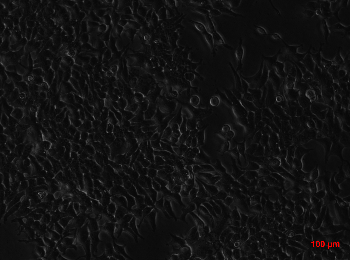

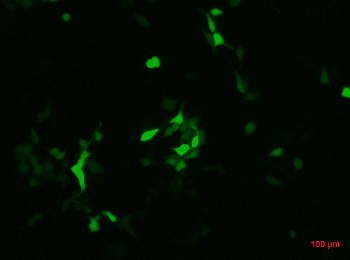

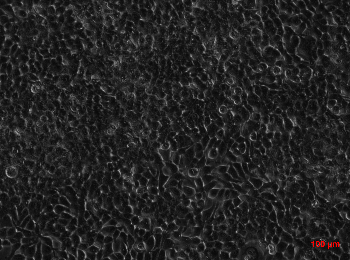

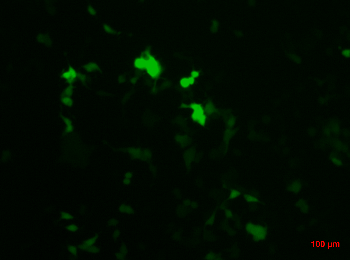

**CSedac5**

**CSedac3**

**Bright field**

**1**

**1**

**1**

**1**

**1**

**1**

**1**

**Fluorescence**

**2**

**2**

**2**

**2**

**2**

**2**

**2**

**Bright field**

**1**

**1**

**1**

**1**

**1**

**1**

**1**

**Fluorescence**

**2**

**2**

**2**

**2**

**2**

**2**

**2**

**A**

**B**

Figure S9 - Representative images of fluorescence microscopy of transfected cells by CSedac3 and CSedac5 polyplexes with 100 µg of IGFBP-3 or -5, respectively. Cells were visualized after 48h and 72h, left and right panel, respectively. Amplification of 100X and scale bar represents 100 µm.
